# A novel mycovirus evokes transcriptional rewiring in the fungus *Malassezia* and stimulates interferon-β production in macrophages

**DOI:** 10.1101/2019.12.18.880518

**Authors:** Shelly Applen Clancey, Fiorella Ruchti, Salomé LeibundGut-Landmann, Joseph Heitman, Giuseppe Ianiri

## Abstract

Mycoviruses infect fungi, and while most persist asymptomatically, there are examples of mycoviruses having both beneficial and detrimental effects on their host. Virus-infected *Saccharomyces* and *Ustilago* strains exhibit a killer phenotype conferring a growth advantage over uninfected strains and other competing yeast species, whereas hypovirus-infected *Cryphonectria parasitica* displays defects in growth, sporulation, and virulence. In this study we identify a dsRNA mycovirus in five *Malassezia* species. Sequence analysis reveals it to be a totivirus with two dsRNA segments: a larger 4.5 kb segment with genes encoding components for viral replication and maintenance, and a smaller 1.4 kb segment encoding a novel protein. Furthermore, RNA-seq of virus-infected versus virus-cured *Malassezia sympodialis* revealed an upregulation of dozens of ribosomal components in the cell, suggesting the virus modifies the transcriptional and translational landscapes of the cell. Given that *Malassezia* is the most abundant fungus on human skin, we assessed the impact of the mycovirus in a murine epicutaneous infection model. Although infection with virus-infected strains was not associated with an increased inflammatory response, we did observe enhanced skin colonization in one of two virus-infected *M. sympodialis* strains. Noteworthy, interferon-β expression was significantly upregulated in bone marrow-derived macrophages when challenged with virus-infected, compared to virus-cured *M. sympodialis*, suggesting that the presence of the virus can induce an immunological response. Although many recent studies have illuminated how widespread mycoviruses are, there are relatively few in-depth studies about their impact on disease caused by the host fungus. We describe here a novel mycovirus in *Malassezia* and its possible implications in pathogenicity.

**Importance:** *Malassezia* species represent the most common fungal inhabitant of the mammalian skin microbiome, and are natural skin commensal flora. However, these fungi are also associated with a variety of clinical skin disorders. Recent studies have reported associations of *Malassezia* with Crohn’s disease and pancreatic cancer, further implicating this fungal genus in inflammatory and neoplastic disease states. Because *M. sympodialis* has lost genes involved in RNAi, we hypothesized *Malassezia* could harbor dsRNA mycoviruses. Indeed, we identified a novel mycovirus of the totivirus family in several *Malassezia* species, and characterized the MsMV1 mycovirus of *M. sympodialis*. We found conditions that lead to curing of the virus, and analyzed isogenic virus-infected/virus-cured strains to determine MsMV1 genetic and pathogenic impacts. MsMV1 induces a strong overexpression of transcription factors and ribosomal genes, while downregulating cellular metabolism. Moreover, MsMV1 induced a significantly higher level of interferon-β expression in cultured macrophages. This study sheds light on the mechanisms of pathogenicity of *Malassezia*, focusing on a previously unidentified novel mycovirus.

## Introduction

Viruses are widely distributed in nature and are found in a variety of hosts, ranging from single-celled organisms to plants and mammals. The presence of mycoviruses, viruses that infect fungi, can be benign but are also associated with toxin secretion in yeast, hypovirulence in pathogenic plant fungi, and hypervirulence in dimorphic fungi (1–5). Mycoviruses are typically composed of double-stranded RNA (dsRNA) genomes, but positive-sense and negative-sense single-stranded RNA viruses, as well as circular ssDNA viruses have been reported to infect fungi (6). Overall, more than 10 different families of viruses have been isolated from fungal hosts (7). Despite similar genomic features shared between mycoviruses and human viruses from the same family, one clear difference between the two is the absence of cell-to-cell transmission proteins in mycoviruses. Therefore, mycoviruses typically lack an extracellular stage and are only transmitted during cell division, or by cell or hyphal fusion (8–10).

A well-known example of a mycovirus is the killer virus of *Saccharomyces cerevisiae*. This toxin-producing dsRNA virus was characterized in the 1970s (11), and found to have two dsRNA segments: a larger, 4.5 kb dsRNA, and a smaller, 1.5 kb satellite dsRNA. The large segment, called L-A, is required for viral replication and maintenance. The smaller segment, called M, encodes a pre-protoxin, which is processed into a mature toxin by host enzymes and then secreted from the cell (12, 13). During the replication cycle of L-A, the single-stranded positive-strand [(+)ssRNA] is transcribed by the virally encoded replication machinery and released from the virion to serve two roles; 1) it is the mRNA template for translation of the Gag and Gag-Pol fusion protein, and 2) it is the template that is packaged into new viral particles where transcription reoccurs (13, 14). Yeast cells harboring both the L-A and M dsRNA segments display a growth advantage over strains lacking these components. When toxin-producing strains compete with wild-type strains, the virus-infected strains secrete the mature killer toxin, killing neighboring cells, whereas the toxin-producing cells remain unharmed (15). A similar fitness advantage is conferred by toxin-producing strains of *Ustilago maydis* (2).

In contrast to the killer viruses that provide virus-infected strains a growth advantage, the mycovirus that infects *Cryphonectria parasitica*, the causative agent of chestnut blight (which decimated the population of chestnut trees in the United States in the early 1900s by killing more than one billion trees) impairs growth of the fungus (16, 17). The growth defect of virus-infected strains results in reduced virulence, hence the name *<u>C</u>ryphonectria <u>p</u>arasitica* <u>h</u>ypovirus (CHV). Five different viral segments isolated from *C. parasitica* have been shown to confer a high level of hypovirulence, although the level of impact on fungal growth observed in the infected fungal strain appears to be virus strain-dependent (16, 18). A mycovirus infecting the filamentous fungus *Pseudogymnoascus destructans* has also been implicated in the epidemiology of white-nose syndrome, a lethal fungal infection in bats, as bats in North America have experienced 5 million deaths in the past decade, and all *P. destructans* isolates from bats in this region harbor a mycovirus (19). This is compared to European and Asian *P. destructans* isolates, which do not harbor the mycovirus, and the resident bat populations appear to be resistant to the endemic fungus.

RNAi is an important pathway conserved in many organisms and serves to regulate gene expression, suppress transposon movement, and maintain genome integrity. Additionally, a functional RNAi pathway has been shown to suppress mycovirus proliferation (20, 21). The biogenesis of small RNAs (sRNAs) originating from the infecting virus (termed virus-derived small interfering RNAs, or vsiRNAs) is analogous to that of endogenous sRNAs, requiring essential components such as dicer (*DCL*), argonaute (*AGO*), and RNA-dependent RNA polymerase (*RDP*) (22). It was previously thought that mycoviruses might only survive in species that lack a functional RNAi pathway as introduction of argonaute and dicer into yeast resulted in the elimination of the mycovirus (20). To this end, fungal species lacking canonical RNAi pathways have been good candidates in which to screen for mycoviruses. However, increased research on mycoviruses over the past decade has produced reports showing that mycoviruses have developed strategies to counteract the RNAi defense mechanisms that suppress antiviral activity (5). In *Aspergillus nidulans*, infection of conidia with Virus 1816 suppresses inverted repeat transgene (IRT)-induced RNA silencing, and when the virus was cleared from ascospores, IRT silencing was restored (23). Other examples of mycovirus-induced RNAi suppression in *C. parasitica* and *Rosellinia necatrix* involve viral targeting of the RNAi machinery. In the case of *C. parasitica*, CHV1-EP713 encodes a papain-like protease (p29) that suppresses hairpin RNA-triggered silencing in the fungus (24). In *R. necatrix*, the mycoreovirus RnMyRV3 s10 gene encodes a product that appears to suppress RNA silencing by interfering with the dicing of dsRNA (25). Most recently, a novel partitivirus, TmPV1, was identified in *Talaromyces marneffei,* and is correlated with the downregulation of the *dcl*-1, *dcl*-2, and *qde*-2 RNAi genes in both virus-infected yeast and mycelia (4). Examples of virus-induced RNAi suppression spanning multiple classes of fungi is evidence that RNAi functions to control viral levels and that mycoviruses have evolved ways to counteract this regulation.

The genus *Malassezia* is a monophyletic group of lipophilic yeasts that colonize sebaceous skin sites and represent more than 90% of the skin mycobiome. Despite being a commensal yeast, *Malassezia* is associated with a number of common skin disorders, such as seborrheic dermatitis, atopic eczema, pityriasis versicolor, and also systemic infections (26). Recently, two landmark studies reported the association of *Malassezia* with gastrointestinal disorders such as Crohn’s disease in a subset of patients with CARD9 mutations (which enhance sensitivity to fungal pathogen-associated molecular patterns [PAMPs]) and pancreatic cancer (27, 28), highlighting the importance of this understudied fungal genus as a crucial component of the human mycobiome. All *Malassezia* species lack the fatty acid synthase gene and thus are lipid-dependent, relying on exogenous fatty acids for growth. This feature makes *Malassezia* unique and fastidious to study under laboratory conditions, and as consequence relatively limited research has been done on the basic biology of this organism.

To date, 18 different species of *Malassezia* have been identified and sequence for 15 of these genomes is publicly available. Of these, the *M. sympodialis* ATCC42132 genome is the most well annotated reference genome based on whole genome sequencing, RNA-seq, and proteomics (29, 30). As commensal inhabitants of the skin, *Malassezia* yeasts interact with the skin immune system, and specific T cells, as well as IgG and IgM antibodies, can be detected in healthy individuals, whereas specific IgE antibodies are prevalent in a large proportion of patients suffering from atopic eczema. A number of molecules with immunoglobulin E (IgE)-binding capabilities have been identified in extracts of *Malassezia* and have been named MalaS allergens. Ten allergens have been identified in *M. sympodialis*, and three in *M. furfur*. Eight of these allergens are conserved proteins that share high similarity with the corresponding mammalian homolog, suggesting that the induction of autoreactive T cells by *Malassezia* allergens could play a role in sustained inflammation (26, 31). Nevertheless, both the fungal and host factors contributing to the transition of *Malassezia* from commensal to pathogen are unknown and this is an active area of research.

*Malassezia* has a relatively small genome of 7.7 Mb, and due to its reduced size has lost nearly 800 genes compared to other sequenced Basidiomycota, including genes of the RNAi pathway (29). Given that *Malassezia* has lost the RNAi machinery, we hypothesized it could harbor mycoviruses and screened our collection of *Malassezia* strains for novel dsRNA viruses. Here, we report the discovery of a novel mycovirus from *M. sympodialis*, MsMV1. Following the screening of three dozen *M. sympodialis* isolates in our collection, we observed that this mycovirus is present in approximately 25% of the isolates. Fungal cells can be cured of the mycovirus upon exposure to high temperature or the antimicrobial agent biphenyl. RNA sequencing of infected and cured strain pairs revealed the complete genome of MsMV1 and illuminated a large shift in the cellular transcriptional profile comparing the congenic virus-infected and virus-cured strains. The presence of MsMV1 correlated with an increased ability to colonize skin in one of two cases tested, and increased production of interferon-β in cultured macrophages was associated with all virus-infected strains tested. These findings reveal a novel mycovirus harbored by *Malassezia* with possible implications for interactions with the host in both the commensal and disease states.

## Results

### Identification and characterization of a novel mycovirus in *Malassezia*

Whole genome sequencing revealed that the *Malassezia* species have lost the genes required for a functional RNAi pathway (29); therefore, we hypothesized the fungus had the capacity to harbor mycoviruses. Following the screening of *M. sympodialis*, *M. pachydermatis,* and *M. furfur* laboratory freezer stocks, dsRNA species were detected in 11 of 36 *M. sympodialis* isolates (30.5%), one of three *M. pachydermatis* isolates (33.3%), and no *M. furfur* isolates. dsRNA species were also found in one of two *M. globosa* isolates, one *M. obtusa* isolate, and one *M. yamatoensis* isolate (Figure 1A, 1B, and Figure S1). For *M. sympodialis*, two dsRNA segments were visualized on an ethidium bromide-stained agarose gels, one at ~4.5 kb and another at ~1.4 kb. The sizes of these dsRNA segments are similar to those found in *S. cerevisiae* strains that produce killer toxins (1), suggesting the segments may correspond to a helper virus and a satellite RNA encoding a toxin precursor. *M. obtusa* and *M. yamatoensis* appear to only have the larger ~4.5 kb species as no dsRNA species of ~1.4 kb was detected. *M. globosa* also has a ~4.5 kb dsRNA species as well as two smaller species of ~500 bp and ~600 bp (Figure 1B).

**Figure 1.**
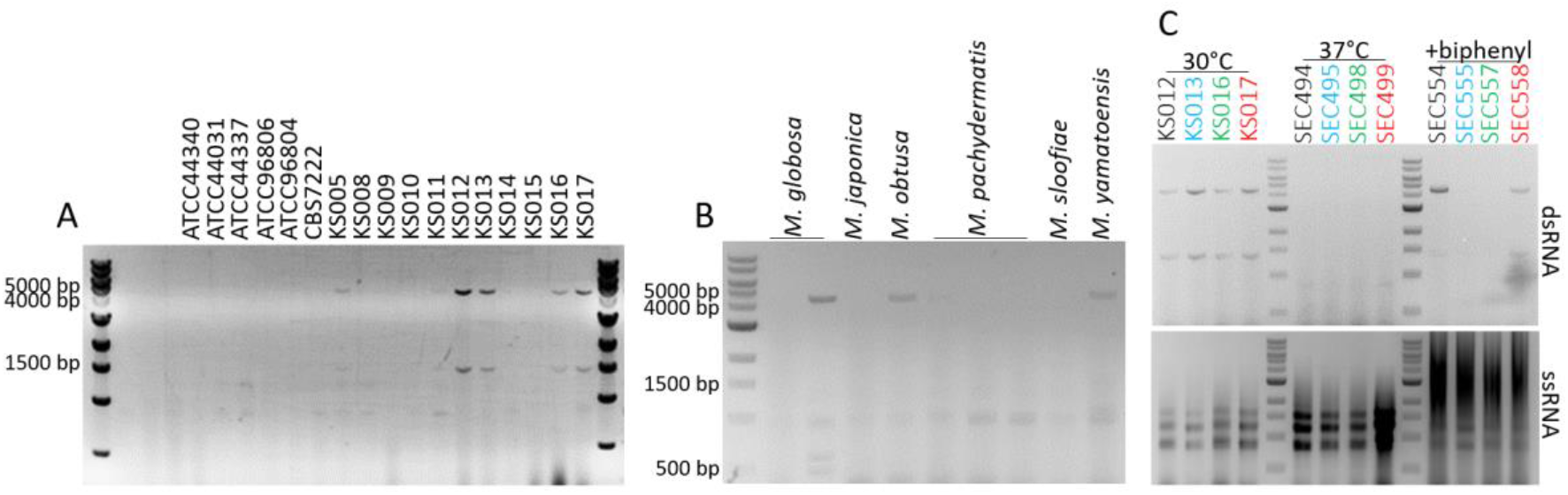
*Malassezia* species harbor dsRNA segments. A) dsRNA from representative *M. sympodialis* strains. B) dsRNA from additional *Malassezia* species. Strain names can be found in Table 1. C) dsRNA (upper panel) and ssRNA (lower panel) from *M. sympodialis* strains passaged at 30°C, 37°C, and on mDixon media containing 10 μg/mL biphenyl. All dsRNA and ssRNA extracts were visualized by electrophoresis in 0.8% agarose gels with 0.5 μg/mL ethidium bromide.

In the present study we focused on the characterization of the mycovirus of *M. sympodialis* as a representative species of the *Malassezia* genus. To unveil the biological and physiological roles of the putative *Malassezia* mycovirus, a crucial first step was to define conditions that allow elimination (i.e. curing) of the mycovirus. In other fungi, mycoviruses can be cured by exposure to stressful conditions or certain drugs. In *S. cerevisiae*, exposure to the fungicide cycloheximide cures killer cells of the killer phenotype (32). This has varying rates of success in other organisms, such as *Aspergillus*, where a less than 50% success rate of mycovirus curing was observed after exposure to cycloheximide (33). In filamentous fungi, conidial isolation and hyphal tipping are more commonly used methods for curing (33), and polyethylene glycol was shown to cure *P. destructans* of PdPV-pa infection (19). To determine if *M. sympodialis* could be cured of the dsRNA, the strains were passaged on modified Dixon’s (mDixon) media and subjected to various stresses. We found that six passages on mDixon medium with no additives at high temperature (37°C) resulted in a complete loss of both dsRNA segments in four *M. sympodialis* virus-infected strains (Figure 1C, middle section). Following a single passage on mDixon medium with 10 μg/mL biphenyl, strains SEC555 and SEC557 derived from KS013 and KS016, respectively, lost both dsRNA species, while strains SEC554 and SEC558 (derived from KS012 and KS017, respectively) retained both dsRNA species (Figure 1C, right section). All dsRNA containing strains retained both segments when passaged on mDixon with no additives at 30°C (Figure 1C, left section).

To determine if the dsRNA moieties isolated from *M. sympodialis* are of viral origin, the dsRNA molecules of *M. sympodialis* were converted to cDNA and subjected to TA-cloning and Sanger sequencing. Clones of the large moiety were obtained from strain KS013, and from KS012 for the smaller one. Primer walking with Sanger sequencing revealed only partial insertions of the two into the cloning plasmid: a 1.3 kb region of the large ~4.5 kb segment, and a 374 bp portion of the ~1.4 kb segment. Blastx of the 1.3 kb sequence derived from the larger dsRNA revealed similarities with other mycoviruses deposited in GenBank, while a blastx search of 374 bp sequence derived from the smaller dsRNA found no hit (see later in the text for details). In parallel with the cDNA synthesis, cloning and sequencing, the congenic virus-infected and virus-cured strains KS012 and SEC494 of *M. sympodialis* were subjected to ribosomal RNA-depleted RNA sequencing. The rationale of the RNA-seq analysis was two-fold; first, allowing a more facile identification of the entire viral genome and second, elucidating possible transcriptomic changes evoked by the mycovirus in the host cells.

For the identification of the viral genome, adaptor-free RNAseq reads derived from *M. sympodialis* virus-infected and virus-cured isolates were assembled with Trinity. The resulting transcripts were searched using the sequences previously obtained through TA-cloning as a query, and two perfect hits (*E* value 0.0) were obtained following a blastn search against the virus-infected strain KS012. Both hits are part of two larger independent contigs (DN8321 of 4613 bp, and DN4986 of 4601 bp; see File S1 legend for details) that subsequent analyses (described in detail below) reveal represent the entire viral genome. The two identified contigs are identical with the exception of a few nucleotides located at the sequence extremities and outside coding regions, likely being residual adaptor or assembly artifacts (File S1). Conversely, blastn analysis of the 1.3 kb viral TA-cloned sequence against the transcriptome of the virus-cured isolate SEC494 found only a small 308 bp hit (*E* value 1e-143) that is part of the *M. sympodialis* ATCC42132 *MRP4* ortholog, which encodes a mitochondrial ribosomal protein of the small subunit (MSYG_4110, accession SHO79760). Combined with nucleic acid analysis based on agarose-ethidium bromide stained gels, this sequencing further supports that the dsRNA had been completely eliminated in the virus-cured strain.

Moreover, PCR analyses with primers designed to specific regions of the large and small dsRNA segments failed to produce amplicons on *M. sympodialis* KS012 genomic DNA, suggesting that the viral sequence is not integrated in the genome but is rather cytoplasmic. This finding is corroborated by *in silico* BLAST searches that failed to detect either large or small segment viral sequences in the *de novo* genome assembly of *M. sympodialis* KS012 (described below).

Using the ExPASy Translate web tool, the 4.6 kb segment was found to contain two open reading frames (ORFs), distinguished by a −1 ribosomal frameshift, a common feature of totiviruses (Figure 2A) (34). A BLASTp search of the ORF produced by a frame 1 translation revealed the RNA-dependent RNA polymerase gene (*RDP*) from *Scheffersomyces segobiensis* virus L (SsV L) with 52.69% homology. A BLASTp search of the ORF produced by a frame 2 translation identified the capsid protein from the same *S. segobiensis* virus, in this case with 53.87% homology (35). Phylogenetic analysis of the Rdp1 sequence obtained from RNA-seq grouped it with Rdp1 sequences from other mycoviruses of the totiviridae family (Figure 2B). Moreover, bidirectional blast analysis using the *S. segobiensis* SsV L virus genome as a query against the *M. sympodialis* KS012 transcriptome assembly found the contigs DN8321 and DN4986 (File S1). These results strongly support that the large dsRNA segment isolated from *M. sympodialis* is a novel mycovirus and encodes proteins required for mycoviral replication and maintenance.

**Figure 2.**
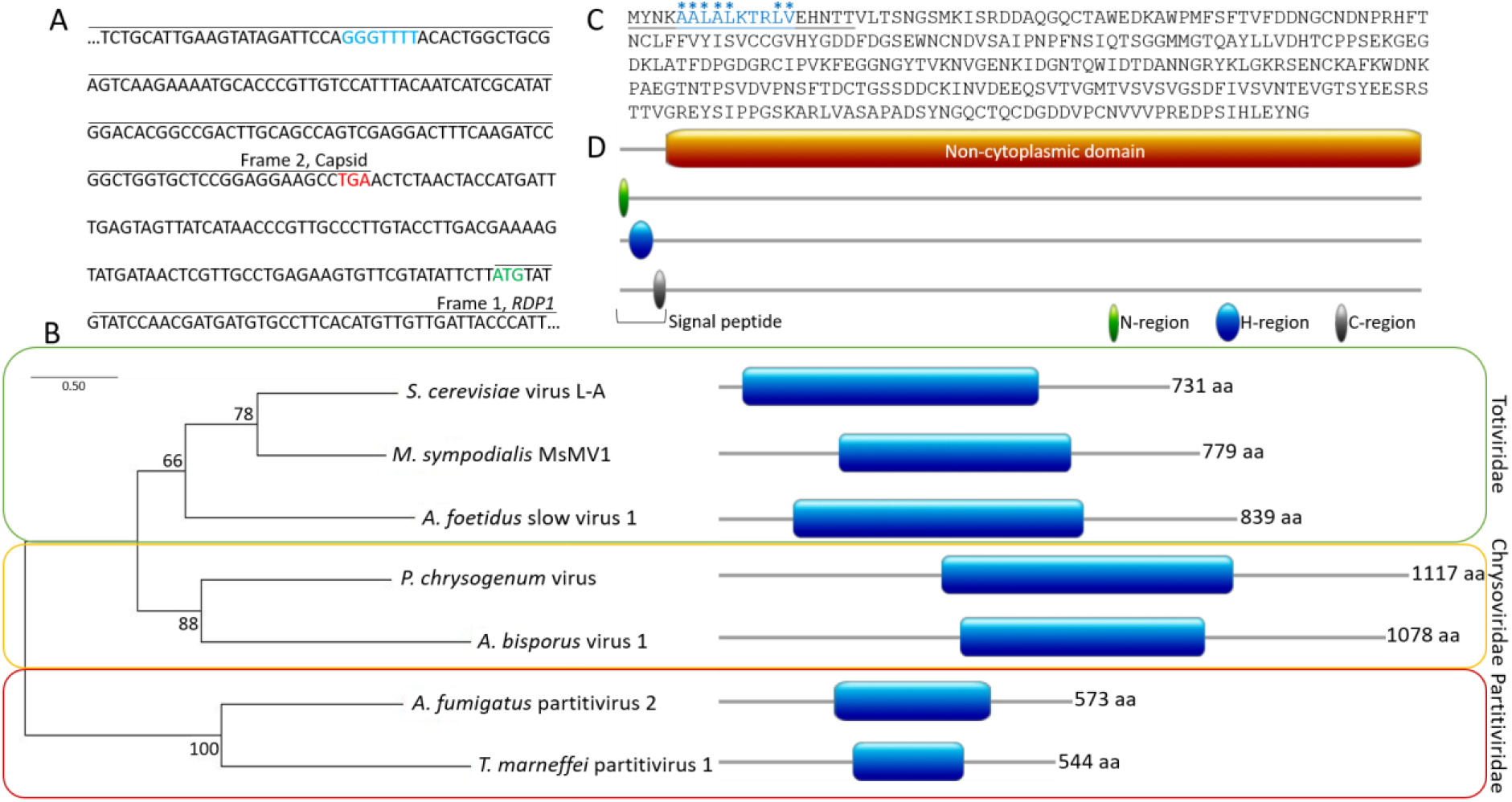
dsRNA segments code for viral protein products. A) Representative region of the large dsRNA segment illustrating the two open reading frames for the capsid and Rdp1 and the site of the −1 ribosomal frameshift. Stop codon for capsid highlighted in red. Start codon for Rdp1 highlighted in green. Frameshift site highlighted in blue. B) Phylogenetic analysis of mycovirus Rdp1 sequences. Scale bar represents the number of substitutions per site. Blue boxes on protein diagrams represent the reverse transcriptase (RT) domain of the Rdp1 protein. C) Protein sequence of the small dsRNA segment. The signal peptide is underlined and hydrophobic residues indicated with stars. D) Pictorial representation of predicted domains of the small dsRNA segment.

Similarly, the 374 bp query sequence from the smaller dsRNA segment identified via TA-cloning was subjected to a blastn search against the assembled transcriptomes of the virus-infected and virus-cured *M. sympodialis* strains KS012 and SEC494, respectively. Two highly significant hits (*E* value 3e −170 and 2e −166) were obtained only against the genome of the virus-infected strain KS012. The two hits are part of two independent contigs (DN7179 of 1546 bp and DN6052 of 1479 bp; see File S2 legend for details), consistent with the size of the small dsRNA segment based on agarose gel electrophoresis. These contigs are identical with the exception of a few nucleotides located at the sequence extremities and outside coding regions (File S2). Translation of this contig showed a single ORF (Figure 2C), which when used in a BLASTp search in NCBI returned only weak (≤29% homology, *E* value ≥1.6) hits to the 3’ region of uncharacterized and hypothetical proteins in *Sporisorium reilianum* and *Streptomyces*, respectively; no significant hits were obtained from NCBI for the small segment.

To infer the function of this virally-encoded protein, the predicted amino acid sequence was subjected to domain prediction using Interpro Scan. A 305 amino acid non-cytoplasmic domain was identified at the C-terminal end of the protein and a short 19 amino acid signal peptide is predicted at the N-terminus (Figure 2D). Signal peptides are commonly present at the N-terminus of proteins that are destined for secretion. Signal peptides are composed of three regions: an N-region, an H-region, and a C-region. Typically, the N-region contains a stretch of positively charged amino acids, the H-region is the core of the signal peptide that contains a stretch of hydrophobic amino acids, and the C-region contains a stretch of amino acids that are recognized and cleaved by signal peptidases (36). The presence of this signal peptide, and the lack of homology to any protein in the NCBI database, suggests the ~1.5 kb dsRNA segment from *M. sympodialis* encodes a previously unidentified, novel protein of unknown function that is likely secreted from the cell. Given the data supporting that these dsRNA segments represent a dsRNA virus in *M. sympodialis*, we termed this virus MsMV1 (*<u>M</u>alassezia <u>s</u>ympodialis* <u>m</u>yco<u>v</u>irus1).

### MsMV1 infection causes an upregulation of a large number of ribosomal genes

As described, total RNA from congenic virus-infected and virus-cured strain pairs KS012 and SEC494 was subjected to analysis of differentially expressed genes in the virus-infected strain compared to its cured counterpart. For RNA-seq analysis, three biological replicates of each sample were analyzed. RNA-seq reads of virus-infected and virus-cured strains KS012 and SEC494 were mapped against the reference *M. sympodialis* genome ATCC42132 using HiSat2. Transcript abundance was estimated through StringTie, and differentially expressed genes with a false discovery rate (FDR) < 0.05 and a log_2_-fold change +/- 1 were identified with EdgeR (37, 38).

The virus-infected strain displayed differential expression of 172 genes in comparison to the virus-cured strain, with 89 genes being upregulated (log_2_-fold change between 1 and 8.14) and 83 genes being downregulated (log_2_-fold change between −1 and −4.6) (Figure 3A). Among the proteins differentially upregulated, ~50% are involved in ribosomal biogenesis and translation (Figure 3B). The remaining genes are involved in DNA replication and maintenance, transcription, stress responses, and other cellular metabolic processes. The most highly upregulated gene in the virus-infected strain, with a log_2_-fold change of 8.14, was MSYG_3592, encodes an uncharacterized mediator complex protein. The mediator complex is a transcriptional coactivator that interacts with transcription factors and RNA polymerase II, which transcribes all protein-coding and most non-coding RNA genes (39). The change in expression of this gene, combined with the upregulation of phosphorylation activity, ribosomal biogenesis, and cytoplasmic translation, indicates a large transcriptional shift in the virus-infected strain.

**Figure 3.**
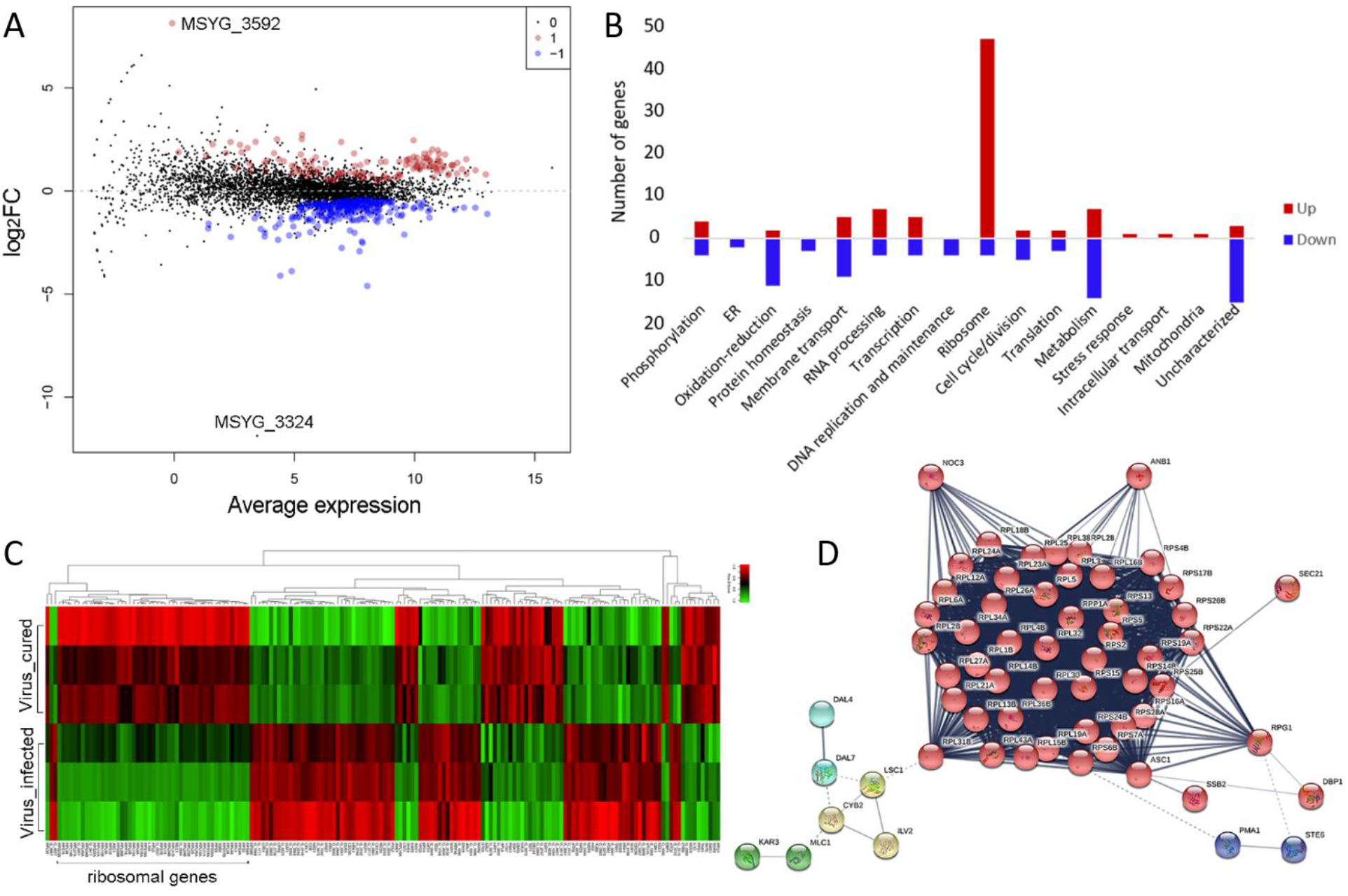
RNA-seq reveals transcriptional rewiring in virus-infected *M. sympodialis* strain. A) Dot plot of log_2_ fold-changes in gene expression in the virus-infected strain KS012. Red and blue dots represent upregulated and downregulated genes that pass a false discovery rate (FDR) of 5%, respectively. Gray dots indicate genes that do not meet the FDR. The virus-cured strain SEC494 was used as the baseline for gene expression, and a gene is marked as differentially expressed in the virus-infected strain KS012 if expression differs from SEC494. B) Number of genes upregulated (red) and downregulated (blue) categorized by gene ontology. C) Heat map of the three RNA-seq technical replicates of virus-infected and virus-cured strains. Green represents upregulated genes and red represents downregulated genes. D) Gene network map of upregulated genes in virus-infected strain. Genes are clustered by color based on the Markov-Cluster (MCL) inflation parameter.

Downregulated genes included those involved in metabolic processes, oxidation-reduction processes, transmembrane transport, cell cycle and cell division, and many uncharacterized transcripts. In particular, the most downregulated gene that passed the FDR <0.05 threshold with a log_2_-fold change of −4.6 is an endoplasmic reticulum (ER) membrane protein, MSYG_1988, a gene whose predicted product is involved in the movement of substances across the ER membrane and/or protein synthesis. The second most downregulated gene is MSYG_0220 (log_2_-fold change = −4.1), which is an ortholog of *S. cerevisiae CDC39.* Cdc39 is a component of the Ccr4-Not1 complex that, in addition to its role as a global transcriptional regulator, regulates mRNA decay. These data suggest that elimination of aberrant mRNA transcripts may be inhibited in the virus-infected strain. Interestingly, we identified a gene (MSYG_3324) with the lowest log_2_-fold change (−11.88) that did not pass the statistically significant threshold of FDR <0.05. The gene MSYG_3324 gene encodes a hypothetical protein that has a guanine nucleotide exchange factor domain, leucine-rich repeats of the ribonuclease inhibitor (RI)-like subfamily, and an N-terminal Herpes BLLF1 superfamily domain, which consists of the BLLF1 viral late glycoprotein that is the most abundantly expressed glycoprotein in the viral envelope of the Herpes viruses. Strikingly, BLLF1 represents a major antigen responsible for production of neutralizing viral antibodies *in vivo* (40), and we speculate that it might be strongly downregulated by the MsMV1 mycovirus as a mechanism to dampen *M. sympodialis* antiviral defense systems. Overall, our RNA-seq dataset is clearly indicative of a significant transcriptional rewiring of *M. sympodialis* in the presence of the dsRNA mycovirus MsMV1.

The three technical replicates of each strain produced highly similar expression profiles, which are distinct from one another (Figure 3C). The largest cluster of upregulated genes in the virus-infected strains included ribosomal components, and a gene network map shows the tight interaction of these ribosomal components (Figure 3D). The large, central cluster consists of ribosomal proteins that are regulated by translation initiation factors Anb1 and Rpg1, and also Noc3, which bind ribosomal precursors to mediate their maturation and intranuclear transport.

One question that we sought to answer was whether the conditions employed to cure the virus changed the strain in any way other than virus elimination. We generated whole genome Illumina data and then performed genome-wide analyses and SNP calling of the *M. sympodialis* cured strain SEC494, comparing it to the virus-infected strain KS012. Only two nucleotide changes unique to the virus-cured isolate SEC494 were identified. These include mutations in the coding regions of *KAE1*, an ATPase, and *CRM1*, a karyopherin responsible for the transport of molecules between the cytoplasm and the nucleus (Table S1, File S3). Being as these mutations were only found in the virus-cured strain, they may be a consequence of the “curing treatment” and we predict their impact on fitness to be minimal given similar growth characteristics of the two strains (Table S1; File S3). Moreover, the assembled KS012 and SEC494 genomes are co-linear with the reference *M. sympodialis* genome ATCC42132 (Figure S2A-C), and no change in ploidy or aneuploidy was observed for any chromosome (Figure S2D).

### *Malassezia* mycovirus does not impact pathogenicity in an epicutaneous murine model

In a recent study, Sparber and colleagues described a murine epicutaneous infection model that involves application of *Malassezia* to the mouse dorsal ear following mild tape stripping (41). Colonization of host tissue (fungal load) is determined by plating homogenized tissue onto agar plates and counting colony forming units; an increase in ear thickness is used as a measurement for skin inflammation. When mice were challenged with congenic virus-infected and virus-cured strains in this model, only the virus-infected strain KS012 displayed increased colonization compared to its virus-cured counterpart, while there was no difference in the other strain pairs tested by day 4 post-infection (Figure S3A). No change in ear thickening (as a readout for inflammation) was observed in response to the virus-infected strains compared to their cured counterparts. To control for the temperature exposure in the virus-cured strains, a strain without MsMV1 was passaged at 37°C. No changes in skin colonization or ear thickening was observed in the heat-treated virus-uninfected counterpart when tested in the epicutancous infection model (Figure S3A). To determine if there was a difference in the immune response elicited by the virus-infected and virus-cured strains, we assessed the expression of interleukin-17 (IL-17), a key mediator of the host response against *Malassezia* on the skin (41). However, we could not detect any statistically significant differences in cytokine induction in response to virus-infected and virus-cured strains (Figure S3C), indicating that IL-17 expression is not regulated by the virus and not responsible for the observed differences in fungal load observed for strain KS012. We also assessed expression of IFN-β, a central player in antiviral immunity, but could not measure expression above the basal levels detected in the skin of naïve control animals (Figure S3C). The same was also true at earlier or later time points after infection (Figure S3D, E). Similarly, the heat-untreated and heat-treated virus-uninfected control strain did not display differential IL-17 or IFN-β expression (Figure S3C).

### Virus-infected strains induce interferon-β expression in bone marrow-derived macrophages in a TLR3-independent manner

Given its close relationship with the skin, *Malassezia* primarily interacts with keratinocytes and tissue-resident mononuclear phagocytes, including macrophages. Therefore, we assessed the interaction of *M. sympodialis* virus-infected and virus-cured strains with macrophages. Bone marrow-derived macrophages (BMDMs) are mature, differentiated bone marrow monocyte/macrophage progenitors that are suitable for host-pathogen interaction studies (42). Interferon-β (IFN-β) expression in BMDMs was measured by RT-qPCR following incubation with virus-infected and virus-cured *M. sympodialis* strains for 24 hours. The presence of MsMV1 induced significantly more IFN-β expression in BMDMs than virus-cured strains (Figure 4A).

**Figure 4.**
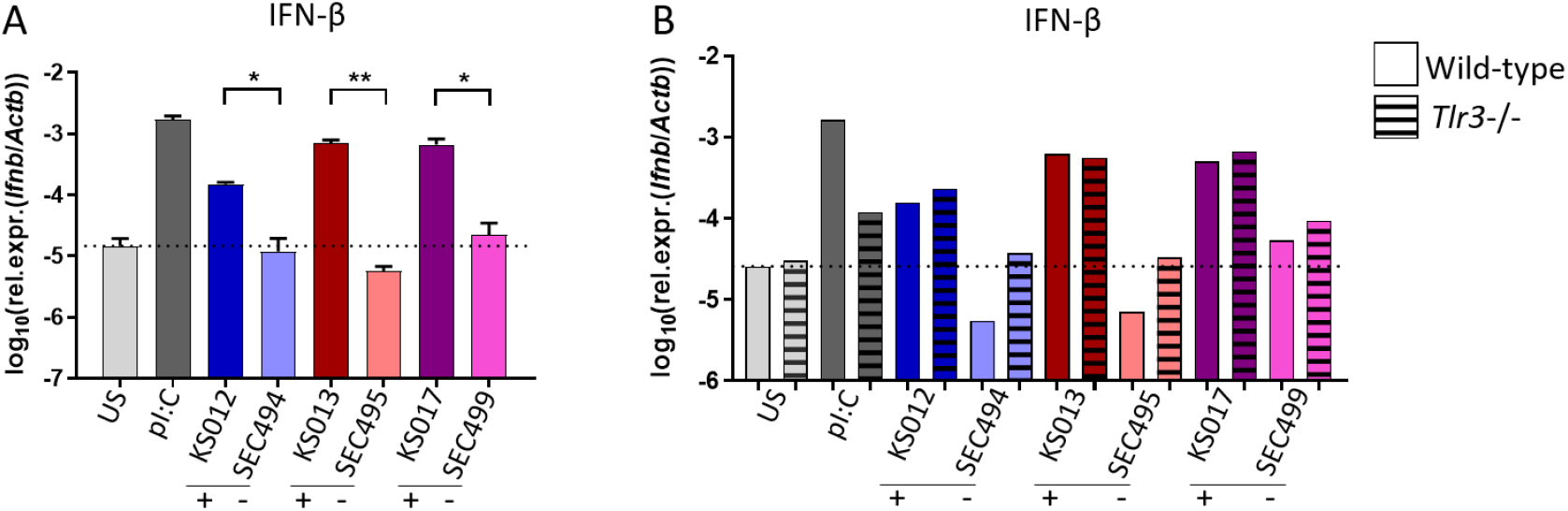
MsMV1 dsRNA virus induces interferon-β (IFN-β) expression in cultured macrophages in a TLR3-independent manner. A) IFN-β expression in bone marrow-derived macrophages (BMDM) after infection with the indicated virus-infected and virus-cured strains at MOI=5 for 24 hours. Bars are the mean + SEM of four replicate samples with each being the pool of two separate wells. Data are pooled from two independent experiments. B) IFN-β expression by wildtype (solid bars) and *Tlr3*−/− (dashed bars) BMDM after infection with the indicated virus-infected and virus-cured strains at MOI=5 for 24 h. Bars are a pool of two wells. Data are representative of one out of two independent experiments. polyI:C (pI:C) was included as a positive control; US, unstimulated. The dotted line indicates the expression levels under unstimulated conditions. Statistics were calculated using unpaired Student’s t test. *p<0.05, **p<0.01. Black bars below the x-axis labels group virus-infected/virus-cured strain pairs. ‘+’ indicates virus-infected, ‘-‘ indicates virus-cured.

In leishmaniasis, a parasitic disease caused by *Leishmania*, an elevated burden of *Leishmania* RNA virus-1 (LRV1) in the parasite is associated with parasite dissemination and increased disease severity (43, 44). Toll-like receptors (TLRs) are innate immune-recognition receptors known to play a role in viral infection, and TLR3, which recognizes and responds to dsRNA (45), was found to be required for the induction of pro-inflammatory factors and disease exacerbation in leishmaniasis (43). Given the relationship between TLR3 and dsRNA recognition, TLR3 deficient BMDMs were incubated with the virus-infected and virus-cured *Malassezia* strains. In contrast to observations with LRV1, the increased production of IFN-β by MsMV1-infected strains was not dependent upon TLR3, as deletion of this gene in BMDMs did not result in a loss of IFN-β expression by the virus-infected strains (Figure 4B).

BMDM viability and macrophage activation, as measured by CD86^+^ surface expression and TNF expression, was also assessed (Figure S4A). No significant differences were found in BMDMs incubated with virus-infected and virus-cured *M. sympodialis* strains, and deletion of TLR3 did not change this result (Figure S4C). The seemingly enhanced induction of CD86 by the virus-cured strain SEC494 (Figure S4A) was not reproducible in independent repeat experiments, not paralleled by other readouts of macrophage activation, and not consistent with the results obtained with other virus-infected and virus-cured strain pairs. Therefore, it seems unlikely to be related to the absence of the virus.

## Discussion

Mycoviruses are proving to be much more ubiquitous in nature than previously appreciated. Fungi harboring mycoviruses display a range of phenotypes, which in some cases can prove beneficial or detrimental for the host plant or animal. For instance, *C. parasitica* carries the CHV1 virus that causes reduced growth and pathogenicity of the fungus. Therefore, trees infected with virus-infected strains of *C. parasitica* display smaller cankers than trees infected with virus-free strains (16). Application of hypovirus-infected strains to cankers has proven to be an effective biocontrol agent as it halted the decimation of the chestnut tree in North America (17). Conversely, virus-infected *L. guyanesis* exacerbates leshmaniasis in mice and increases parasitemia, leading to the dissemination and metastasis of the parasite from the infection site (43).

Here, we describe the discovery of dsRNA mycoviruses in five *Malassezia* species: *M. sympodialis, M. pachydermatis, M. globosa, M. obtusa,* and *M. yamatoensis* (Figure 1A, 1B, and Figure S1), and characterize in detail the MsMV1 virus from *M. sympodialis.* RNA-sequencing revealed the large dsRNA segment from *M. sympodialis* is of viral origin and contains two open reading frames that encode the Rdp1 and capsid proteins, which are translated by a −1 ribosomal frameshift, a hallmark feature of the Totiviridae virus family (Figure 2A)(34). Accordingly, phylogenetic analysis of the Rdp1 open reading frame groups most closely with other mycoviruses from the totiviridae family (Figure 2B). Using the small dsRNA segment sequence as a query, no significant matches were found in the NCBI database, suggesting the satellite virus is unique to this fungus and could serve a specialized role. Taken together, these data support the discovery of a novel mycovirus in the important skin fungus *M. sympodialis*. Further investigation will be required to characterize the presumptive mycoviruses in the other *Malassezia* species.

We find that exposure to high temperature (37°C) or to an antimicrobial agent (biphenyl) cures *M. sympodialis* of MsMV1 (Figure 1C). These are two conditions frequently encountered by fungi. Systemic human patient fungal isolates are exposed to high temperature during infection, and isolates from both patients and the environment are often purified on media containing antimicrobials to inhibit bacterial or filamentous fungal growth from the host or environment. It is possible these typical microbiological isolation methods may be inducing mycovirus curing during isolation. Taken together with the growing reports on their discovery of novel mycoviruses, mycoviruses may be much more abundant than currently appreciated. Not all of the *Malassezia* isolates analyzed in this study harbored a mycovirus, and this could reflect either natural diversity or curing during isolation. It is not clear why these two conditions cause virus curing. In the case of high temperature, this stressful condition may alter cellular physiology rendering the cell a less favorable host for the mycovirus. Viral replication and maintenance, while neutral at 30°C, could interfere with cellular processes required for survival at 37°C. Moreover, high temperature is known to increase heat shock and misfolded proteins in the cell (46), and if full Hsp90 function is required for viral replication then the virus could be lost due to the overburdened Hsp90.

RNA-sequencing of virus-infected and virus-cured strains revealed major differences in the transcriptional profile between the two strains. Most notably is the upregulation of a large number of ribosomal components in the virus-infected strains, suggesting an increased translational activity induced by the mycovirus. A similar result was found with the plant fungus *Sclerotinia sclerotiorum*, which harbors the SsHV2-L hypovirus, where rRNA metabolic processes and cellular component biogenesis were the classes of the most upregulated genes in virus-infected strains (47). Alternatively, the opposite has been observed in *S. cerevisiae*, where virus-infection is associated with the down-regulation of ribosome biogenesis and stress response, suggesting that in *S. cerevisiae*, virus-infected cells may be less stressed due to host protection systems for the killer virus (48).

Genes involved in oxidation-reduction processes are another category that was downregulated in the virus-infected strain. Denitrification, the biological process of converting nitrate to ammonia through reduction, which is then broken down into nitrogen, nitrous oxide, and nitric oxide (NO), is an important pathway in living organisms. NO is an important signaling molecule in bacteria and fungi. Furthermore, NO production is positively correlated with aflatoxin production in *A. nidulans*, where deletion of flavohemoglobin, a nitric oxide reductase, leads to reduced sterigmatocystin production and *aflR* expression (49). Flavohemoglobin, encoded for by the *YHB1* gene in *M. sympodialis*, does not appear to play a role in putative toxin producing strains as it is downregulated only 0.69-fold in MsMV1-infected strains. Recent work from Wisecaver and colleagues as well as our lab has identified that *Malassezia* acquired flavohemoglobin via a horizontal gene transfer event from Actinobacteria (49, 50). A likely explanation for the difference in the role of flavohemoglobin between *A. nidulans* is because the gene was not endogenous to *Malassezia* and rather acquired from an exogenous bacterial source, and therefore may not have been adapted to play a regulatory role with the virus. The presence of the virus may be inducing the upregulation of transcriptional and translational machinery in order to synthesize components required for viral replication, maintenance, and persistence. Analysis of gene expression in additional virus-infected strains as well as virus-uninfected strains would further confirm the gene expression changes imparted by the virus. How this signal is translated from the virus to the fungus is an interesting area for future research.

RNAi is a common antiviral defense system that operates in fungi to combat viral persistence in the cell. As expected, *C. parasitica* CHV1-infected cells exhibit an upregulation of silencing-related genes in an effort to reduce the viral burden or clear the virus. Alternatively, a different phenomenon occurs in *T. marneffei* TmPV1-infected cells where the virus can subsequently dampen expression of silencing-related genes and blunt the cellular RNAi defense mechanism (4, 51, 52). *M. sympodialis* has lost the canonical components of RNAi and hypothesize that the loss of RNAi may be contributing to mycovirus replication and persistence. Furthermore, we observed changes in genes related to regulating mRNA processing. One significantly downregulated gene is involved in mRNA decay, MSYG_0220 (*CDC39*). As described earlier, during the virus replication cycle, a ssRNA intermediate is formed prior to virion assembly (13) and is vulnerable to fungal defense mechanisms. Suppressing the expression of genes involved in eliminating anomalous mRNA transcripts could be a strategy for the virus to survive in the fungal cell. Interestingly, the RNA-seq analysis identified the downregulation of MSYG_03961 (*CPR3*) in virus-infected *M. sympodialis* cells. *CPR3* is a gene that encodes a mitochondrial cyclophilin (53) that could be linked to pathogenesis via established roles of mitochondria in basidiomycete pathogens (54). Although the passing at high temperature did not induce many genomic changes, it is possible epigenetic changes occurred during the treatment that resulted in changes in gene expression, and further studies are needed to address this question.

Mycoviruses impact fungal virulence in different ways. Mycoviruses can be asymptomatic, whereas others induce either a hypo- or hyper-virulent state in the host fungus. We employed a recently developed murine model for *Malassezia* in which the fungus is applied onto the mouse ear following mild barrier disruption to replicate hallmarks of atopic dermatitis (41). We found one of two virus-infected strains that displayed an enhanced ability to colonize the murine skin, compared to its virus-cured counterpart, but did not observed any differences in inflammation or cytokine production (Figure S3A, B, and C). Promisingly, cultured macrophages responded differently to virus-infected and virus-cured strains in which more IFN-β was produced from BMDMs incubated with virus-infected strains than virus-cured strains (Figure 4A). Other parameters such as BMDM viability and BMDM activation were measured in parallel and no difference was detected between virus-infected and virus-cured strains (Figure S4A). We conclude that under certain circumstances, MsMV1 appears to provide an enhanced ability to colonize skin, and increased IFN-β in cultured macrophages is a common characteristic of all virus-infected strains tested, thus suggesting some role in the pathogenicity of *M. sympodialis.* However, the degree to which the virus plays a role is still uncertain. It is possible characteristics in the genomic background of the *M. sympodialis* virus-infected strain influences the degree to which the virus exacerbates disease. Perhaps there are features of the KS013 genome that allow it to better control viral load of MsMV1 and so it does not impart the same growth advantage on the skin. As KS012 was the only virus-infected genome sequenced in this study we do not know if there are any differences between it and KS013 that would account for the differences observed in skin colonization experiments.

Due to the role of IFN-β in the stimulation of antiviral and antiproliferative products (55), it is possible that the production of IFN-β may lead to enhanced clearing of virus-infected fungal cells from macrophages. TLR3 binds dsRNA in mammalian cells and was shown to play a critical role in disease exacerbation in leishmaniasis when the infecting *Leishmania* strain harbors a mycovirus (43, 44). In the case of *M. sympodialis*, we did not observe a requirement of TLR3 for IFN-β production in response to virus-infected strains (Figure 4B). We posit that dsRNA recognition by *Malassezia* may occur through an alternative virus recognition pathway yet to be elucidated that may recognize mycoviral dsRNA as a fungal pathogen-associated molecular pattern (PAMP).

In conclusion, we report for the first time evidence of mycoviruses infecting the fungal genus *Malassezia*. Although the target of the putative satellite toxin is unknown, it induces an immune response in macrophages. Co-incubation of the virus-infected *M. sympodialis* with virus-cured *M. sympodialis* and bacterial species (*Escherichia coli, Corynebacterium* spp., and *Staphylococcus* spp.) failed to reveal antifungal or antibacterial properties of the virus-infected *M. sympodialis* strains. Determining if changes in the expression of metabolic genes in the virus-infected strains results in resistance to or susceptibility to certain environment conditions would be interesting to investigate in the future. More research will be required to determine if this possible mycovirus toxin plays a role in common skin disorders caused by *Malassezia* by targeting the host, bacteria, or fungi of the skin microbiome.

## Materials and methods

### Strains

*Malassezia sympodialis* strains were stored at −80°C in 25% glycerol stocks and maintained on modified Dixon’s (mDixon) medium at 30°C. mDixon media is composed of 3.6% malt extract, 1% mycological peptone, 1% desiccated ox bile, 1% Tween 60, 0.4% glycerol, and 2% agar for solid medium.

### RNA extraction and dsRNA enrichment

Total RNA was extracted from *M. sympodialis* liquid cultures as follows. 5 mL liquid mDixon cultures were grown at 30°C until saturation, approximately 3-4 days. Cultures were pelleted at 3,000 rpm in a table top centrifuge for 3 minutes, washed with 5 mL dH_2_O, and pellets were frozen at −80°C until lyophilization overnight. Following lyophilization, pellets were broken up with 3 mm glass beads and the RNA extraction was performed using TRIzol (ThermoFisher, 15596026) according to the manufacturer’s instructions. RNA was resuspended in 50 μl diethyl pyrocarbonate (DEPC) water. RNA was treated with TURBO DNase (ThermoFisher, #AM2238) according to manufacturer’s instructions, and resuspended in 50 μl DEPC water. ssRNA precipitation/dsRNA enrichment was done overnight at −20°C by combining 25 μl of DNase treated RNA, with 28 μl 8 M LiCl ([final]=2.8 M) and 27 μl DEPC water for a final volume of 80 μl. The mixture was centrifuged at 4°C for 10 minutes and the supernatant was recovered and transferred to a fresh eppendorf tube (the pellet can be saved and resuspended in 25 μl DEPC water as a ssRNA control). To the supernatant, 7.5 μl 3 M NaCl ([final]=0.1 M) and 187.5ul 100% ethanol ([final]=85%) was added and placed at −20°C for at least 1 hour. dsRNA was pelleted at 4°C for 15 minutes and resuspended in 25 μl DEPC water. To visualize dsRNA and ssRNA bands, 10 μl was electrophoresed on an 0.8% agarose gel stained with 5 μg/mL of ethidium bromide.

### TA cloning of dsRNA segments

Large and small dsRNA segments were purified from *M. sympodialis* strains KS012, KS013, KS016, and KS017 as described above. Large and small dsRNA segments were excised from the agarose gel and purified via gel extraction using the QIAquick Gel Extraction Kit (Qiagen, 28704) 3’ adapter sequences from the Illumina TruSeq Small RNA Library Preparation Kit (Illumina, RS-200) were ligated onto the dsRNA species by mixing together 1 μl of adapter with 1 μg dsRNA in a 6 μl reaction and heated at 70°C for 2 minutes. Following heating, tubes were immediately placed on ice. To the adapter reaction, 2 μl of HML ligation buffer, 1 μl RNasin, and 1 μl T4 RNA Ligase 2 (truncated) was added and mixed by pipetting. The solution was incubated for 1 hour at 28°C.

cDNA synthesis was achieved by combining 5 μl of adapter-ligated dsRNA with 1 μl RT primer and heated at 70°C for 2 minutes, then immediately placed on ice. Next, the adapter-ligated/RT primer mixture was used as template with the AffinityScript cDNA Synthesis Kit (Agilent, 600559) with the following reaction conditions: 25°C for 5 minutes, 55°C for 15 minutes, 95°C for 5 minutes. cDNA was PCR amplified using the RT primer, cloned into the pCR-Blunt vector using the Zero Blunt PCR Cloning Kit (Invitrogen, K270020), and transformed into *E. coli* with One Shot TOP10 Chemically Competent *E. coli* (Invitrogen, C404003). Plasmids were extracted from purified bacterial colonies and subjected to Sanger sequencing (Genewiz) using universal M13F and M13R primers.

### RNA sequencing

5 mL liquid mDixon cultures were grown at 30°C for 3 days. Cells were pelleted at 3,000 rpm in a table top centrifuge for 3 minutes, washed with 5 mL dH_2_O, and resuspended in 1 mL dH_2_O. The 1 mL cell suspension was transferred to a bead beating tube with acid washed 425-600 μm beads. Cells were pelleted with beads in a microcentrifuge at 6,000 rpm for 2 minutes. The supernatant was removed and the cell pellet was flash frozen in liquid nitrogen. Frozen bead/cell pellets were stored at −80°C until RNA extraction. To prepare RNA, bead/cells pellets were thawed on ice and 1 mL TRIzol (ThermoFisher, 15596026) was added. Cells were bead beated 10 times in 1 minute ‘on’, 1 minute ‘rest’ cycles. Following bead beating, 200 μl of chloroform was added and the RNA extraction continued as described in the manufacturer’s instructions. Final RNA pellets were treated with TURBO DNase (ThermoFisher, AM2238) according to manufacturer’s instructions, and RNA was resuspended in 50 μl of nuclease free water (Promega, P119A). Illumina 150 bp PE libraries were prepared using the TruSeq Stranded Total RNA-seq Kit in combination with the Illumina Ribozero Yeast Kit. Library preparation and RNA sequencing was performed at the Duke University Center for Genomic and Computation Biology.

After sequencing, Illumina raw reads were trimmed with Trimmomatic to remove Illumina adaptors (56). For the identification of the dsRNA mycovirus genome, one sequencing replicate of both the virus-infected and virus-cured strains was assembled into a transcriptome using Trinity (57), which was run on Galaxy (58). The generated transcriptome was searched using the TA-cloned sequences derived from both the large and small dsRNA segments. *In silico* functional characterization of the mycovirus sequences was performed through blast analyses and InterPro scan (https://www.ebi.ac.uk/interpro/search/sequence/). For RNA-seq analysis, the most recent *M. sympodialis* strain ATCC42132 genome assembly and annotation were used as a reference (30). Illumina reads from three independent biological replicates were trimmed with Trimmomatic as reported above, and cleaned reads were mapped to the *M. sympodialis* reference genome using HiSat. Generated .bam files were used to run StringTie with the *M. sympodialis* annotation as guide, and the −e option to provide the correct output for the statistical analysis (37). Read count information for statistical analysis were extracted using a provided python script (prepDE.py). EdgeR was used to determine the differentially expressed genes (DEGs) in the *M. sympodialis* virus-infected strain KS012 compared to its cured counterpart SEC494. StringTie and EdgeR were run on Galaxy (58). DEGs were considered those with a FDR < 0.05 and with a log_2_-fold change ±1. Functional annotation of the DEGs was performed using the Blast2GO pipeline, which includes the BLASTx against the NCBI non-redundant protein database, gene ontology (GO) annotation and InterProScan (59). Heatmap for the DEGs was generated using the web-based server heatmapper with the average linkage for hierarchical clustering (http://www.heatmapper.ca/expression/), and gene network interaction were determined using STRING (https://string-db.org/cgi/input.pl) with the *S. cerevisiae* protein set as reference.

### Whole genome DNA sequencing and assembly

25 mL liquid mDixon cultures of *M. sympodialis* KS012 and SEC494 were grown at 30°C for 3 days. Cells were pelleted at 3,000 rpm in a table top centrifuge for 3 minutes, washed with 10 mL dH_2_O, and cell pellets were frozen at −80°C. Cells pellets were lyophilized overnight, pulverized using 3 mm glass beads, and subjected to genomic DNA extraction using the CTAB method (60). Library preparation and 50 bp PE Illumina sequencing was performed at the Duke University Center for Genomic and Computation Biology. Illumina reads for *M. sympodialis* KS012 and SEC494 were assembled using SPAdes with default parameters (61). Dot plot comparisons were carried out using nucmer with the –nosymplify, --filter, and –fat parameters; nucmer was run on Galaxy. The *de novo* genome assembly of *M. sympodialis* KS012 was used for blast searches using both large and small dsRNA viral sequences.

Illumina reads of the *M. sympodialis* virus-infected strain KS012 and virus-cured SEC494 were mapped to the *M. sympodialis* reference ATCC42132 using BWA (62), and the resulting .sam file converted to .bam with samtools (63). Bam files were converted in .tdf format using IGV, and the read coverage for *M. sympodialis* virus-infected strain KS012 and virus-cured SEC494 visualized in the IGV browser using the *M. sympodialis* ATCC42132 as reference. For variant analysis, the .bam files were generated as above and the *M. sympodialis* reference ATCC42132 genome were used to run samtools mpileup; variant analysis was carried out using the bcftools call option; the resulting .vcf file was merged with the *M. sympodialis* annotation using the variant effector predictor software present on the EnsemblFungi database (https://fungi.ensembl.org/Malassezia_sympodialis_atcc_42132_gca_900149145/Info/Index).

### Phylogenetic analysis

The MsMV1 Rdp1 was aligned with representative viral Rdp1 sequences found on NCBI using the Mega7 software. Phylogenies were built using 100 bootstrap values using the Maximum Likelihood method based on the Tamura-Nei model (64). Protein domains were predicted using NCBI Blastp and images were created using the MyDomain Image Creator from PROSITE on the ExPASy Bioinformatics resource portal.

### Murine topical infection model

WT C57BL/6j mice were purchased by Janvier Elevage and maintained at the Laboratory Animal Science Center of University of Zurich, Zurich, Switzerland. Mice were used at 6-12 weeks in sex- and age-matched groups. Epicutaneous infection of the mouse ear skin was performed as described (41, 65). Briefly, *Malassezia* strains were grown for 3-4 days at 30°C, 180 rpm in liquid mDixon medium. Cells were washed in PBS and suspended in native olive oil at a density of 20 OD_A600_/mL. 100 μl suspension (corresponding to 2 OD_A600_) of yeast cells was applied topically onto the dorsal ear skin that was previously disrupted by mild tape stripping while mice were anaesthetized. Ear thickness was monitored prior and after infection using an Oditest S0247 0-5 mm measurement device (KROEPLIN) and the increase was calculated by subtracting the values measured prior to the experiments from those measured at the endpoint. For each animal, the mean of the ear thickness from both ears was calculated. For determining the fungal loads in the skin, tissue was transferred in water supplemented with 0.05% Nonidet P40 (AxonLab), homogenized and plated on modified Dixon agar and incubated at 30°C for 3-4 days.

### Macrophage infection *in vitro*

Macrophages were generated from WT C57Bl/6j, *Tlr3*^*−/−*^ (66) (Swiss Immunological Mouse Repository (SwImMR) hosted by the University of Zurich) bone marrow cells in the presence of 20% L929 conditioned R10* medium. After 6 days of culture, cells were detached using PBS/5% FCS/2.5 mM EDTA and seeded at a density of 10^5^ cells/100 μl in 96-well tissue culture plates (for transcript analysis) and 10^6^ cells/1 mL in 24-well plates (for FACS analysis). Cells were rested overnight and infected the next day with *Malassezia* at the indicated MOI for 16 hours (for FACS analysis) or for 24 hours (for transcript analysis). PolyI:C (50 μg/mL) was included as a control.

### Flow cytometry

Macrophages were gently detached by using cell scrapers and stained with antibodies directed against CD11b (clone M1/70), F4/80 (clone BM8), CD86 (clone GL-1), and MHC-II (clone M5/114.15.2) in PBS supplemented with 1% FCS, 5 mM EDTA and 0.02% NaN_3_. LIVE/DEAD Near IR stain (Life Technologies) was used for exclusion of dead cells. Cells were acquired on a FACS Cytoflex (Beckman Coulter) and the data were analyzed with FlowJo software (FlowJo LLC). The gating of the flow cytometric data was performed according to the guidelines for the use of flow cytometry and cell sorting in immunological studies (67), including pre-gating on viable and single cells for analysis.

### RNA extraction and quantitative RT-PCR

Isolation of total RNA from ear skin and from BMDM cultures was carried out according to standard protocols using TRI Reagent (Sigma Aldrich). cDNA was generated by RevertAid reverse transcriptase (ThermoFisher). Quantitative PCR was performed using SYBR Green (Roche) and a QuantStudio 7 Flex (Life Technologies) instrument. The primers used for qPCR were as follows: *Il17a* forward 5’-GCTCCAGAAGGCCCTCAGA-3’, reverse 5’-AGCTTTCCCTCCGCATTGA-3’ (68); *Tnf* forward 5’-CATCTTCTCAAAATTCGAGTGACAA-3’, reverse 5’-TGGGAGTAGACAAGGTACAACCC-3’ (69); *Ifnb* forward 5’-AACCTCACCTACAGGGC-3’, reverse 5’-CATTCTGGAGCATCTCTTGG-3’ (43); *Actb* forward 5--CCCTGAAGTACCCCATTGAAC-3’, reverse 5’-CTTTTCACGGTTGGCCTTAG-3’. All RT-qPCR assays were performed in duplicate and the relative expression (rel. expr.) of each gene was determined after normalization to *Actb* transcript levels.

## Data availability

Both sequencing reads for *M. sympodialis* virus-infected strain KS012 and *M. sympodialis* virus-cured strain SEC494 are available in the NCBI SRA database (BioProject PRJNA593722). The large and small dsRNA fragments of the mycovirus genome have been deposited in GenBank under accession numbers MN812428 and MN831678, respectively.

## Acknowledgements

We thank Marco A. Coelho for performing the genomic assemblies of the virus-infected and virus-cured *M. sympodialis* KS012 and SEC494 strains. This work was supported by NIH/NIAID R37 award AI39115-22 and R01 award AI50113-15 awarded to JH and by SNSF grant 310030_189255 awarded to SLL. JH is a Co-Director and Fellow of the CIFAR program Fungal Kingdom: Threats & Opportunities. We thank Blake Billmyre and other members of the Heitman lab, as well as Florian Sparber, Amanda McLeod, and Jessica Shannon for helpful discussions and comments on the manuscript.

---

**Supplemental Figure 1.** *Malassezia* species harbor dsRNA segments. dsRNA from select *M. sympodialis* (A) and *M. furfur* (B) strains was analyzed by electrophoresis on a 0.8% agarose gel with 0.5 μg/mL ethidium bromide.

**Supplemental Figure 2.** Genomic comparisons of *M. sympodialis* virus-infected and virus-cured strains compared to the reference strain ATCC42132. (A-C) Dot plot comparison of the *M. sympodialis* strain ATCC42132 with the virus-infected *M. sympodialis* KS012 (A) and the virus-cured SEC494 (B), and of KS012 and SEC494. (C) Read coverage of the *M. sympodialis* strains KS012 and virus-cured SEC494 compared to the haploid reference strain ATCC42132

**Supplemental Figure 3.** Select virus-infected strains exhibit increased ability to colonize skin in a murine ear skin infection model. A) Fungal load, B) ear thickening, and C) cytokine expression in the skin of mice that were epicutaneously infected with the indicated virus-infected and virus-cured strain pairs for 4 days, D) 2 days, or E) 7 days. The mean + SEM of each group is indicated. Data in A-C are pooled from two independent experiments with n=6 mice for all groups, except n=3 for KS015 and SEC497. DL, detection limit. Statistics were calculated using One-way ANOVA. \**p*<0.05, \*\*\*\**p*<0.0001. Black bars below the x-axis labels group virus-infected/virus-cured strain pairs. ‘+’ indicates virus-infected, ‘-‘ indicates virus-cured.

**Supplemental Figure 4.** Macrophage viability, CD86 expression, and TNF expression is not affected by the presence of MsMV1 in *M. sympodialis* strains. A) BMDM were infected with the indicated virus-infected and virus-cured strains at MOI=1 and analyzed for viability and CD86 surface expression by flow cytometry at 16 hours post-infection or infected at MOI=5 and analyzed for TNF expression by RT-qPCR at 24 hours post-infection. Each bar represents the mean + SEM of four samples, each being the pool of two separate wells. Data are pooled from two independent experiments. The dotted line indicates the expression levels under unstimulated conditions. B) Flow cytometry gating strategy used to assess BMDM viability (top row) and CD86 expression (bottom row). C) BMDM viability, CD86 surface expression and TNF expression of wild-type and *Tlr3*−/− BMDM infected with the indicated virus-infected and virus-cured strains and analyzed as in A). Data are from one out of two representative experiment with each value being the pool of two wells. Statistics were calculated using an unpaired Student’s t test. *p<0.05. ‘+’ superscripts indicate virus-infected strains, ‘-‘ superscripts indicate virus-cured.

**Table 1.** Strains used in dsRNA gel images. Strains are listed top to bottom in order of gel wells from left to right. If known, clinical data and sample location are listed. U = unknown, H = healthy individual, D = dandruff, TV = tinea versicolor, PV = pityriasis versicolor, SD = seborrheic dermatitis, OE = otitis externa. ‘*’ indicates strains sampled from an animal.

**Table 2.** Strains used in this study.

**Table S1.** Variants present only in the *M. sympodialis* virus-cured SEC494 compared to the virus-infected KS012.

**File S1.** Identification of the large dsRNA segment of the *M. sympodialis* MsMV1. Note that the designations “TRINITY_DN8321_c1_g1_i1 len=4613 path=[4591:0-4612] [−1, 4591, −2]” and “TRINITY_DN4986_c1_g1_i1 len=4601 path=[4579:0-4600] [−1, 4579, −2]” are automatically generated by the software Trinity. We arbitrary used the denomination DN8321 and DN4986 to indicate the contigs that represent the *M. sympodialis* MsMV1 mycovirus genome, as identified by blast analyses. Considering “TRINITY_DN8321_c1_g1_i1 len=4613 path=[4591:0-4612] [−1, 4591, −2]”, according to the software instructions, TRINITY_DN8321_c1_g1_i1 indicates Trinity read cluster ‘TRINITY_DN8321_c1’, gene ‘g1’, and isoform ‘i1’. ‘Len’ indicates the length of the cluster (contig) in bp, and path information indicates the path traversed in the Trinity compacted de Bruijn graph to construct that transcript.

**File S2.** Identification of the small dsRNA segment of the *M. sympodialis* MsMV1. For the contigs designation, see explanation in the legend of File S1.

**File S3.** Screenshot of the variants presents in the coding regions of the genes *KAE1* and *CRM1* in the *M. sympodialis* strains SEC494.

