## Supplementary figures and images for "A novel mycovirus evokes transcriptional rewiring in the fungus *Malassezia* and stimulates interferon-β production in macrophages"

### Figure S1

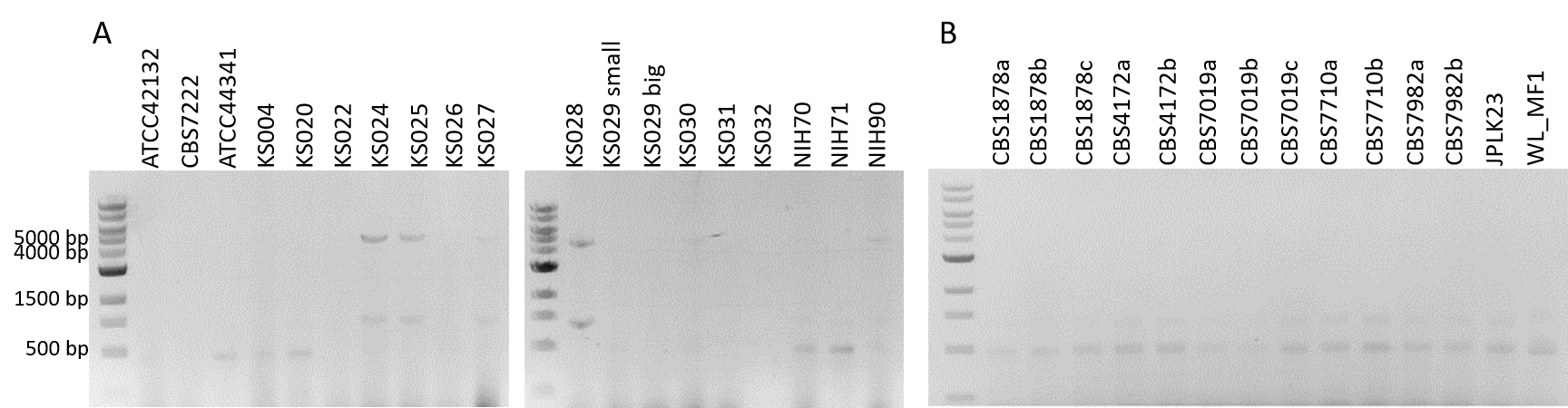

### Figure S2

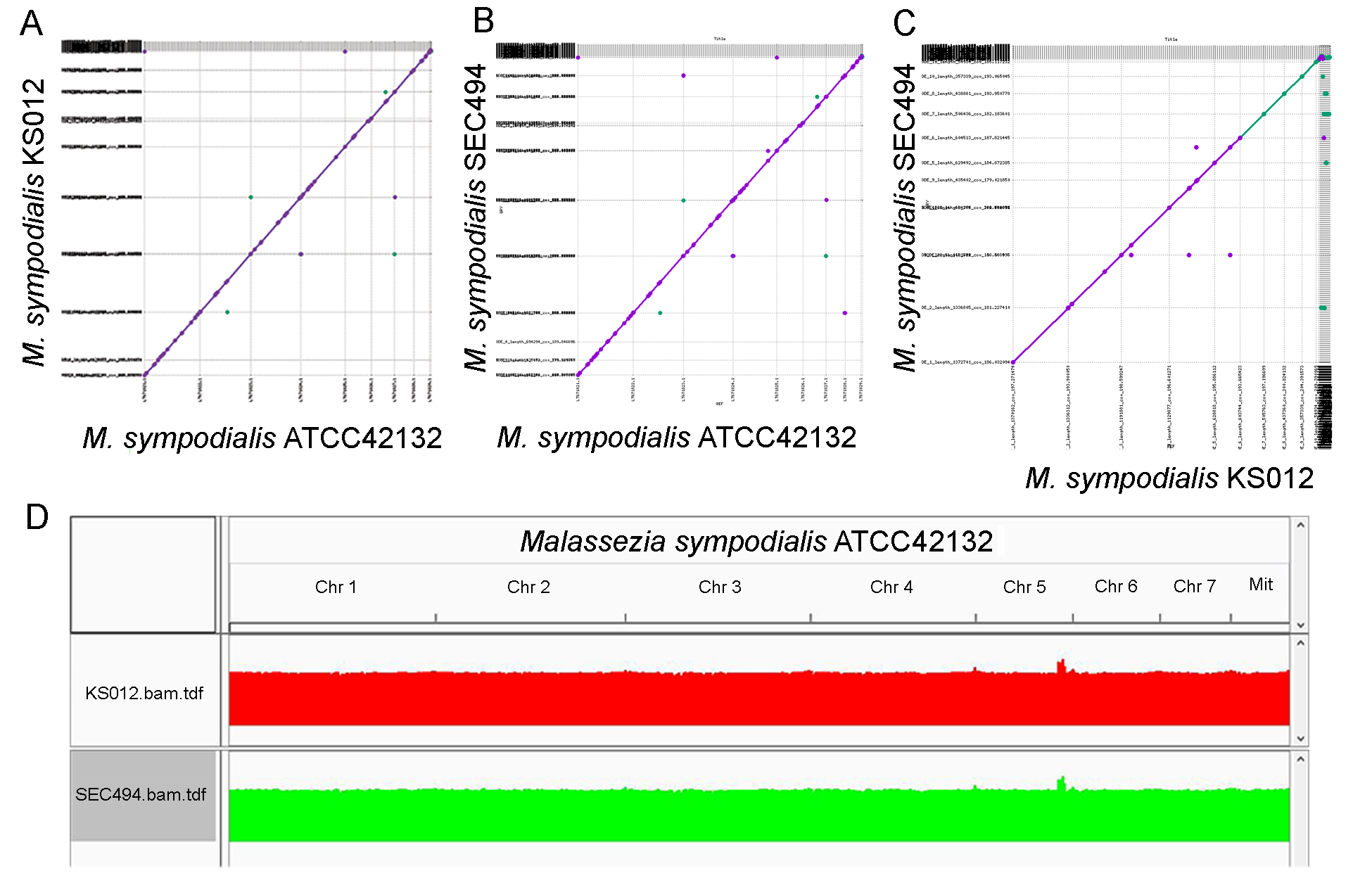

### Figure S3

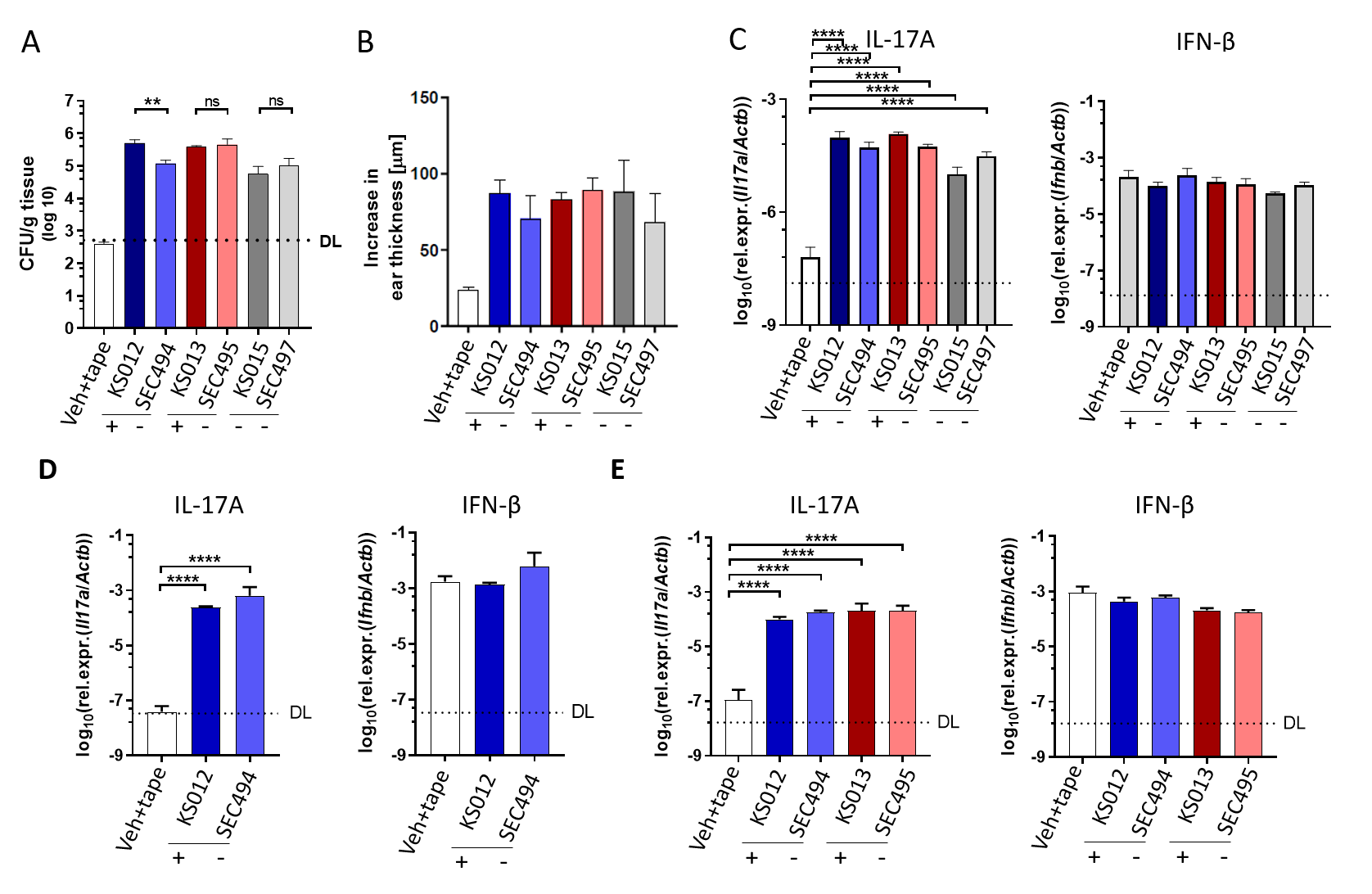

### Figure S4

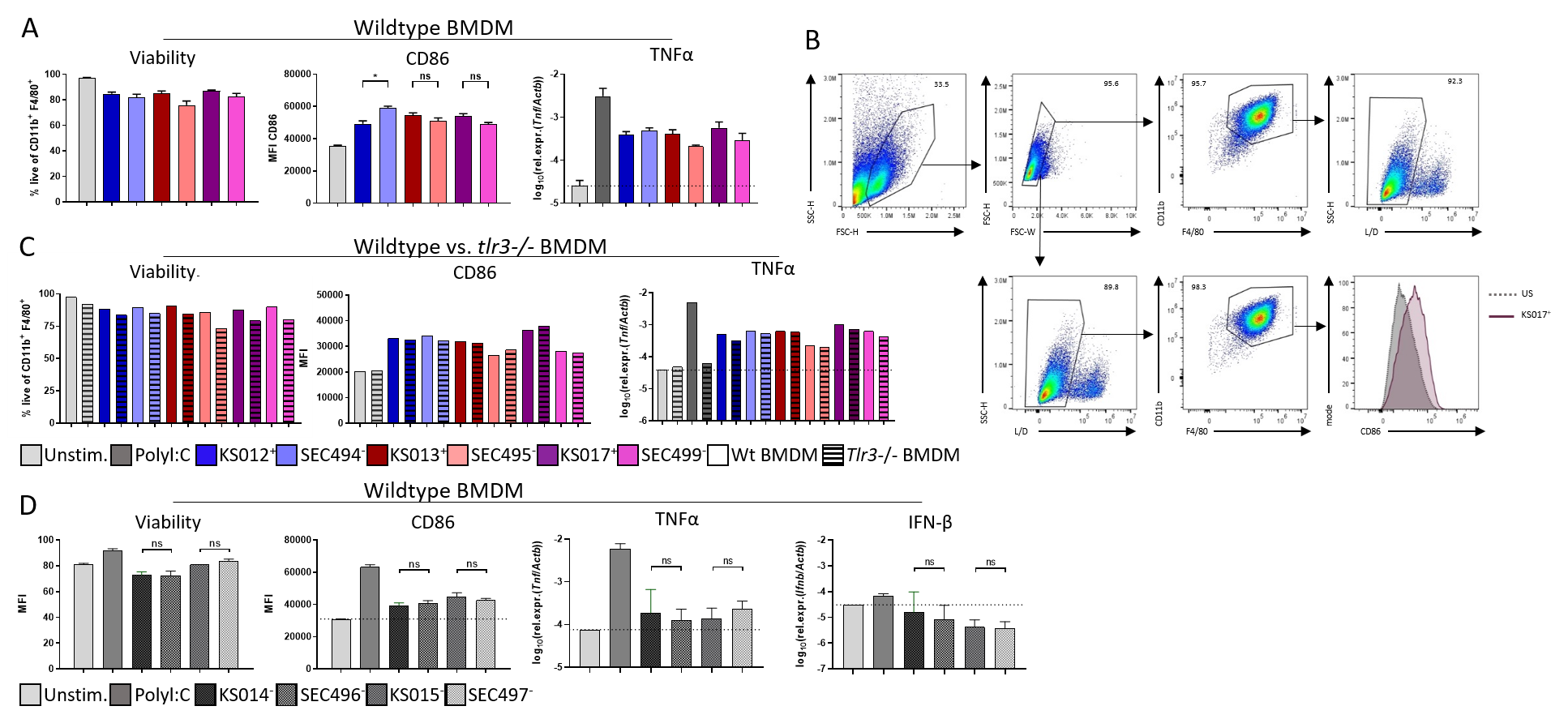
